# Replicative ageing perturbs the metabolic signature of murine C_2_C_12_ skeletal myotubes

**DOI:** 10.1101/2022.04.20.488970

**Authors:** Daniel G. Sadler, Marie M Phelan, Jonathan Barlow, Richard Draijer, Helen Jones, Dick H. J. Thijssen, Claire E. Stewart

## Abstract

**Introduction:** Chronological ageing is associated with mitochondrial dysfunction and increased reactive oxygen species (ROS) production in skeletal muscle. However, the effects of replicative ageing on skeletal muscle cellular metabolism are not well known. Using an established myoblast model of cellular (replicative) ageing, we investigated the impact of ageing on energy metabolism in murine C_2_C_12_ myotubes.

**Methods:** Control (P7-11) and replicatively ‘aged’ (P48-51) C_2_C_12_ myoblasts were differentiated over 72-120 h. Mitochondrial bioenergetics were investigated by respirometry and mitochondrial superoxide and cellular ROS were measured in the absence and presence of antimycin A (AA). Genes related to mitochondrial remodelling and the antioxidant response were quantified by RT-qPCR. Intracellular metabolites were quantified using an untargeted ^1^H-NMR metabolomics pipeline.

**Results:** Mitochondrial coupling efficiency (Control: 79.5 vs. Aged: 70.3%, *P*=0.006) and relative oxidative ATP synthesis (Control: 48.6 vs. Aged: 31.7%, *P*=0.022) were higher in control vs. aged myotubes, but rates of mitochondrial superoxide production were lower (Control: 2.4×10^−5^ ± 0.4 × 10^−5^ vs. Aged: 9.7×10^−5^ ± 1.6×10^−5^ RFU/sec/cell; *P*=0.035). Replicatively aged myotubes had greater mRNA expression of mfn2 and Tfam compared to control. Yet, Nrf2 and PGC-1α expression were 2.8-fold and 3.0-fold higher in control versus aged myotubes over 24 h and 48 h (*P*<0.05), respectively. Branched chain amino acids L-leucine, L-isoleucine and L-valine, and L-carnitine were less abundant in aged versus control myotubes.

**Conclusion(s):** Replicative ageing is associated with bioenergetic uncoupling, increased ROS production and impaired amino acid metabolism. Our findings suggest that cellular mitochondrial dysfunction and altered energy metabolism may exacerbate the age-related decline in skeletal muscle mass and function.

## Introduction

Skeletal muscle is essential for locomotion, maintenance of posture and energy storage. With advancing age, there is a progressive loss of skeletal muscle mass and function, otherwise known as sarcopenia (Cruz-Jentoft et al., 2010). This condition is prevalent in adults over 65 years of age and increases the risk of falls, mobility disability and early mortality (Clark & Manini, 2010). Adult skeletal muscle has a remarkable capacity to regenerate and maintain physiological function, owed to its population of myogenic stem cells, or satellite cells. However, loss of satellite cell function with older age impedes the regenerative capacity of skeletal muscle, thereby exacerbating the gradual loss of muscle mass and functionality (Chen et al., 2020).

The mechanisms underlying sarcopenia are not completely understood, although mitochondrial dysfunction is thought to play a key part (López-Otín et al., 2013). Mitochondrial oxidative capacity (ADP stimulated) was associated with muscle strength and quality in 326 older adults (Zane et al., 2017), whereas mitochondrial coupling, maximal oxidative capacity and hydrogen peroxide emission were strongly related to muscle quality in a cohort of 29 older adults (Distefano et al., 2018). Direct examinations of skeletal muscle mitochondrial respiratory function have revealed chronological ageing to be associated with augmented proton leak (Porter et al., 2015), impaired maximal respiration (Tonkonogi et al., 2003) and blunted ADP sensitivity (Holloway et al., 2018). However, it has become clear that age-related impairments to mitochondrial function may simply be an artefact of lower habitual physical activity levels of older versus younger adults (Distefano et al., 2018; St-Jean-Pelletier et al., 2017). Therefore, chronological ageing *per se* may not impair muscle mitochondrial function.

Cellular (replicative) ageing is distinct from chronological ageing but may also alter the phenotype of human skeletal myoblasts. Replicative ageing was previously reported to attenuate the expression of myogenic regulatory factors critical for human myoblast differentiation (Bigot et al., 2008) - a finding that has been replicated in murine C_2_C_12_ skeletal myoblasts (Brown et al., 2017; Sharples et al., 2011). Interestingly, recent evidence suggests replicative ageing alters human myoblast mitochondrial function. Pääsuke and colleagues reported reduced complex I- and complex II-driven respiration of permeabilized human-derived skeletal myoblasts that underwent 3 passages, regardless of the age of the human donor **(Pääsuke et al**., **2016)**. In a separate study using plate-based respirometry, mitochondrial function was comparable between young and elderly human-derived skeletal myoblasts and myotubes (Marrone et al., 2018), although elderly myotubes presented greater rates of mitochondrial superoxide production compared to young when stressed with N-acetyl cysteine or high glucose concentrations. Overall, research suggests that replicative ageing impairs indices of mitochondrial function, but the impact of replicative ageing on energy metabolism of murine skeletal myotubes is not clear.

Therefore, our objectives were to investigate whether replicative ageing affects mitochondrial function and the metabolic signature of murine C_2_C_12_ myotubes. We hypothesised that replicative ageing would impair indices of mitochondrial respiratory function and lower reliance on oxidative phosphorylation which would be reflected in the metabolic signatures of murine C_2_C_12_ myotubes.

## Methodology

### Cell culture and treatment

Commercially available C_2_C_12_ murine skeletal myoblasts (ATCC; Rockville, USA) at passages 8-12 (referred to as ‘control’) and passages 46-50 (replicatively aged and referred to as ‘aged’, having undergone 130-140 population doublings (Sharples et al., 2011)) were used in this study. Following the plating of cells onto gelatinised (0.2% gelatine) culture-plates in growth medium (GM: Dulbecco’s Modified Eagle Medium (DMEM), 10% heat-inactivated foetal bovine serum (FBS), 10% heat-inactivated new-born calf serum, 2 mM L-glutamine, and 1% penicillin-streptomycin), confluent C_2_C_12_ myoblasts were washed twice with phosphate-buffered saline (PBS, (Sigma-Aldrich, Poole, UK) and switched to pre-warmed (37°C) differentiation medium (DM: DMEM, 2% heat-inactivated horse serum, 2 mM L-glutamine, 1% penicillin-streptomycin). After switching myoblasts to DM, they were allowed to differentiate for 72-96 h.

### Mitochondrial ROS production

Mitochondrial superoxide was detected in C_2_C_12_ myotubes using the targeted MitoSOX probe (Invitrogen, Thermo Fisher Scientific, Waltham, USA). C_2_C_12_ cells were seeded at 3 × 10^4^ cells/mL into 12-well plates, and at ∼80% confluence, were switched to DM. After 96 h differentiation, myotubes were washed 2 × in PBS, and switched into pre-warmed Krebs-Ringer buffer (KRH) comprising: 135 mM NaCl, 3.6 mM KCl, 10 mM HEPES (pH 7.4), 0.5 mM MgCl_2_, 1.5 mM CaCl_2_, 0.5 mM NaH_2_PO_4_, 2 mM glutamine and 25 mM D(+)-glucose, with or without 15 µM antimycin A, an inhibitor of complex III (AA; serving as positive control) and incubated at 37ºC for 30 minutes. Next, AA-containing KRH was removed, and MitoSOX was loaded into cells in fresh pre-warmed KRH to a final concentration of 2.5 µM. Plates were immediately transferred to a CLARIOStar plate reader (BMG Labtech, Bucks, Great Britain), and fluorescence was monitored continuously at 30-sec intervals over 30 min. Fluorescent MitoSOX oxidation products were excited at 510 nm and light emission was detected at 580 nm. The plate reader’s focal height and gain were fixed between experiments. Upon completion of the 30-min reading, plates were immediately fixed in 1% (v/v) acetic acid in methanol for determination of cell density by the Sulforhodamine B (SRB; Sigma-Aldrich, Poole, UK) assay.

### Cellular ROS

Cellular reactive oxygen species (ROS) were detected using the CellROX^®^ Deep Red (Invitrogen, Thermo Fisher Scientific, Waltham, USA) by spectrophotometry. Briefly, C_2_C_12_ myoblasts were seeded at 3 × 10^4^ cells/mL into gelatinised 12-well plates, and at ∼80% confluence, switched to DM. After 96 h differentiation, myotubes were washed 2 × in PBS, and switched into pre-warmed KRH, with or without 15 µM antimycin A (AA) and incubated at 37ºC for 30 minutes. Next, KRH was removed, and CellROX was loaded into myotubes in fresh, pre-warmed KRH buffer, to a final concentration of 2.5 µM. Following 30 minutes incubation with the reagent, cells were washed 2 × with PBS and immediately transferred to a plate reader (BMG Labtech, Bucks, Great Britain), where fluorescent CellROX oxidation products were excited at 640 nm and light emission detected at 665 nm. The plate reader’s focal height was optimised and fixed between experiments. Upon completion of the reading, plates were immediately fixed as above for the determination of cell density by the SRB assay, which was used to normalise fluorescence values.

### Mitochondrial respiration

Mitochondrial respiration was measured in adherent C_2_C_12_ myotubes using a Seahorse XFe96 Analyzer (Agilent, Santa Clara, CA, USA). Control (passages 9-11) and replicatively aged (passages 47-50) C_2_C_12_ myoblasts were seeded in XFe96 well plates (Agilent, Santa Clara, CA, USA) at 10,000 cells per well in 100 µL of GM for 24 h to allow cell attachment. After 24 h, C_2_C_12_ myoblasts were washed twice with PBS and transferred to DM. Myotubes were allowed to form over 120 h in DM, with fresh media replacement at 48 h. Sensor cartridges for the XFe96 Analyzer were hydrated with deionised water at 37°C in a non-CO_2_ incubator in the 24 h preceding the assay. On the day of the assay, C_2_C_12_ myotubes were washed into 200 µL pre-warmed modified Krebs Ringer buffer (KRH: 135 mM NaCl, 3.6 mM KCl, 10 mM HEPES, 0.5 mM MgCl_2_, 1.5 mM CaCl_2_, 0.5 mM NaH_2_PO_4_, 2 mM glutamine and 25 mM D(+)-glucose) at pH 7.4. The cells were incubated in this buffer for 45 minutes at 37°C in a non-CO_2_ incubator and then transferred to a Seahorse XFe96 extracellular flux analyser (maintained at 37°C). After an initial 15-minute calibration, oxygen consumption rates (OCR) were measured by a 3-4 loop cycle consisting of a 1-min mix, 2-min wait and 3-min measure to record cellular respiration. After measuring basal respiration, 2 mM oligomycin was added to selectively inhibit the mitochondrial ATP synthase. Subsequently, 2 µM carbonyl cyanide-4-phenylhydrazone (FCCP) was added to uncouple oxygen consumption from ATP synthesis rates, followed by a mixture of 2 µM rotenone and 2 µM antimycin A to determine non-mitochondrial respiration. Rates of oxygen consumption were corrected for non-mitochondrial respiration and expressed relative to the cell DNA content of the appropriate well, determined by the QuantiT^™^ PicoGreen^®^ (ThermoFisher, Waltham, MA, USA) assay. The raw values of extracellular acidification rate (ECAR) and OCR were divided into component rates to calculate the relative contribution of glycolytic (ATP_glyc_) and oxidative ATP-producing reactions (ATP_ox_) to total ATP production, as previously described (Mookerjee & Brand, 2015). Three independent experiments were performed to assess mitochondrial respiration, each containing four technical replicates.

### RT-qPCR – Gene expression quantification

C_2_C_12_ myotubes were lysed in 250 µL TRIzol and total RNA was extracted using the phenol-chloroform method. RNA concentrations were determined by spectrophotometry (NanoDrop™ 2000, Thermo Fisher Scientific, Waltham, USA). Specific primers used in each polymerase cain reaction (PCR) are outlined in supplementary Table 1. After preparation, reaction tubes were transferred to a Rotor-Gene Q PCR thermal cycler for product amplification using a one-step protocol (QuantiFast SYBR^®^ Green RT-PCR Kit, Qiagen, UK). The amplification protocol was as follows: reverse transcription (10 minutes at 50°C), transcriptase inactivation and initial denaturation (95°C for 5 min) followed by 40 × amplification cycles consisting of: 95°C for 10 s (denaturation) and 60°C for 30 s (annealing and extension); followed by melt curve detection. Critical threshold (C_T_) values were derived from setting a threshold of 0.08 for all genes. To quantify gene expression, C_T_ values were used to quantify relative gene expression using the comparative Delta Delta C_T_ (2^-ΔΔCT^) equation (Livak & Schmittgen, 2001), whereby the expression of the gene of interest was determined relative to the internal reference gene (Rp2ϕ) in the treated sample, compared with the untreated zero-hour (72 h differentiation) control.

### Cell Extraction and preparation for NMR

Myotubes were lysed in 50:50 v/v ice-cold acetonitrile:water (HPLC grade) and sonicated in 3 × 30 s bursts using a micro-tip sonicator (50 kHz) in an ice-bath. Extracts were centrifuged at 4°C, 21,500 × g for 5 min to pellet cell-debris with clarified supernatant lyophilised and stored at −80°C prior to NMR acquisition. Lyophilised cell extracts were resuspended to a final sample composition of 100% ^2^H_2_O with 100 mM sodium phosphate buffer pH 7.4, 100 μM Trime-thylsiylpropionate (TSP) and 0.1% azide.

### NMR Set-Up and Acquisition

Spectra were acquired on Bruker 700 MHz avance IIIHD spectrometer equipped with TCI cryoprobe and chilled autosampler (SampleJet, Ettlingen, Germany). Standard vendor pulse sequences were applied to collect 1D ^1^H-NMR spectra (cpmg1dpr). A Carr–Purcell–Meiboom– Gill (CPMG) edited pulse sequence was employed to attenuate signals from macromolecules present (proteins, etc.). Cell extract spectra were collected at 25 °C with 128 transients for optimal sensitivity and 4s interscan delay. Full parameters sets were deposited along with raw and processed spectra in the EMBL European Bioinformatics Institute (EBI) repository MetaboLights (Haug et al., 2013) with ID MTBLS2024.

### Spectra Processing and Quality Control

All spectra were automatically pre-processed at spectrometer by Fourier-transformation, phase correction, 0.3Hz line broadening and baseline correction using standard vendor routines (apk0.noe) and referenced directly to TSP. Spectra were subjected to quality control criteria as recommended by Metabolomics Standards Initiative (MSI) (Considine & Salek, 2019; Sumner et al., 2007). Quality control criteria consisted of appraisal of baseline, line-width, residual water signal width, phase and signal-to-noise. Spectra were bucketed according to peaks boundaries defined with each bucket, the sum of the integral for that region divided by the region width.

### Metabolite Annotation and Identification

Metabolites were annotated via the use of metabolite recognition software Chenomx (Chenomx v 8.2, Chenomx Ltd, Edmonton, AB, Canada), and the respective buckets were annotated prior to statistical analysis. Metabolite identities were confirmed (where possible) via comparison to the in-house metabolite library. Full metabolite annotation and identities are available with the deposited data at EBI.ac.uk/metabolights/MTBLS2024.

### Statistical analysis

To compare outcome measures between control and aged myotubes, independent t-tests were performed. To determine main effects and related interactions between two independent factors, a two-way ANOVA was performed. When main effects and interactions were present, multiple comparisons were performed using Dunnett’s or Sidak’s test where appropriate. Data are presented as means ± SEM, and significance was accepted when *P*<0.05. NMR metabolomics data was analysed by t-test, principal component analysis (PCA) and partial least squares discriminant analysis (PLS-DA). Metabolomics t-test results were subjected to false discovery rate adjustment (Benjamini-Hochberg) of *P*-values to account for multiple variables analysed. Unsupervised multivariate analysis employed PCA whereas supervised multivariate analysis was performed via cross-validated PLS-DA. PLS-DA analysis used a random sample (70%) to train the model and the remaining 30% to test with receiver operator curve to estimate model quality (Westerhuis et al., 2008). Variable in projection (VIP) scores over 1 were used to inform on metabolites responsible for discrimination between groups.

## Results

### Replicative ageing perturbs mitochondrial function and augments ROS production

Rates of basal respiration (Control: 0.37±0.03 vs. Aged: 0.33±0.0.05 pmol/min^-1^/ng DNA^-^ _1_, *P*=0.543), proton leak (Control: 0.07±0.01 vs. Aged: 0.10±0.02 pmol/min^-1^/ng DNA^-1^, *P*=0.304), and ADP phosphorylation (Control: 0.29±0.02 vs. Aged: 0.23±0.03 pmol/min^-1^/ng DNA^-1^, *P*=0.183) were comparable between control and aged myotubes (see Figure 1B). By contrast, maximal respiration (Control: 0.93±0.07 vs. Aged: 0.35±0.05 pmol/min^-1^/ng DNA^-1^, *P*=0.002), spare respiratory capacity (Control: 0.56±0.06 vs. Aged: 0.04±0.02 pmol/min^-1^/ng DNA^-1^, *P*=0.001) and coupling efficiency (Control: 79.5±1.0% vs. Aged: 70.3±1.4%, *P*=0.006; Figure 1C) were all significantly higher in control vs. aged myotubes, suggesting that replicative ageing impairs mitochondrial function of murine myotubes. Rates of ATP production in control and aged myotubes were also assessed, and the relative contribution of J_ATPglyc_ (Control: 51.4±3.2 vs. Aged: 68.3±3.4 %) and J_ATPox_ (Control: 48.6±3.2 vs. Aged: 31.7±3.4 %) to J_ATPproduction_ were significantly higher and lower in aged vs. control myotubes, respectively (*P*=0.022; Figure 1D).

**Figure 1.**
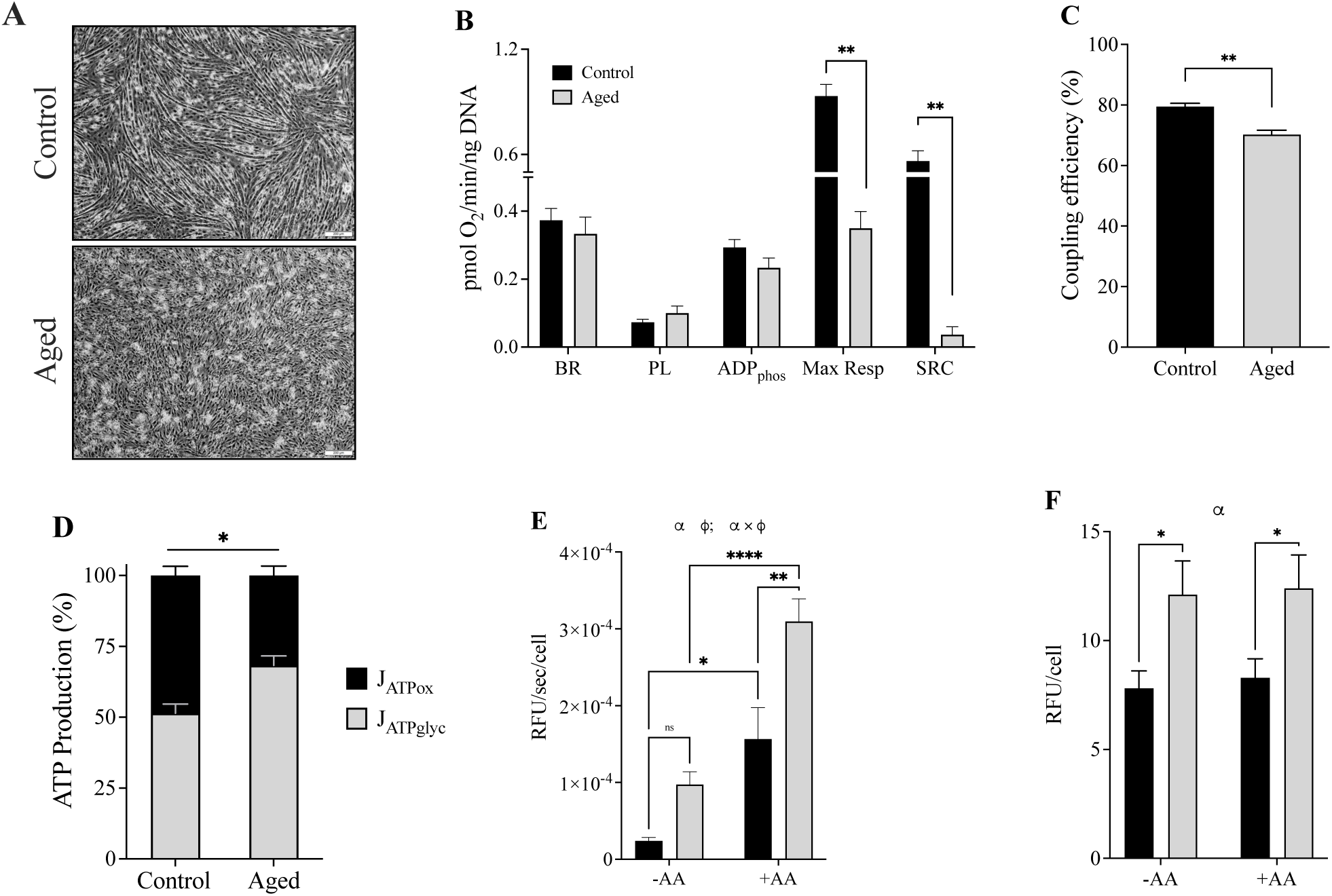
Replicative ageing impairs indices of mitochondrial function. A) Representative images of murine control and aged skeletal myotubes (scale bar = 200 µm). B) Mitochondrial respiration during a mitochondrial stress test. C) Mitochondrial coupling efficiency (%). D) Relative contribution of OXPHOS (black bar) and glycolysis (grey bar) to total ATP synthesis. E) Rates of mitochondrial superoxide production. F) Cellular reactive oxygen species production. Control (black bars) and aged (grey bars) myotubes. BR, basal respiration; PL, proton leak; ADPphos, ADP phosphorylation; Max Resp, maximal respiration; SRC, spare respiratory capacity. Data are mean±SEM, representative of 3 independent experiments and normalised to DNA or cell protein content. α significant main effect of age; ϕ significant main effect if antimycin A. \**P*<0.05, \*\**P*<0.01 and \*\*\*\**P*<0.0001.

Next, rates of mitochondrial ROS production were determined in control and aged myotubes under CTRL conditions (see Figure 1E). A significant main effect of antimycin A (*P*<0.0001) and age (*P*<0.0001) was found on rates of MitoSOX oxidation, and a significant antimycin A × age interaction was observed (*P*=0.031). In the absence of antimycin A, MitoSOX oxidation rates were significantly lower in control compared to aged myotubes (Control: 2.4×10^−5^ ± 0.4 × 10^−5^ vs. Aged: 9.7×10^−5^ ± 1.6×10^−5^ RFU/sec/cell; *P*=0.035). Likewise, control myotubes demonstrated significantly lower rates of MitoSOX oxidation compared to aged when cultured in the presence of antimycin A (Control +AA: 15.7×10^−5^±4.1×10^−5^ vs. Aged +AA: 31.0×10^−5^±2.9×10^−^ _5_ RFU/sec/cell; *P*<0.0001). The impact of replicative ageing on cellular ROS (not mitochondrial specific) was next investigated. Under CTRL conditions, there was a significant main effect of age (*P*=0.003), but not antimycin A (*P*=0.752) on CellROX oxidation. CellROX oxidation was significantly reduced in control compared to aged myotubes (Control: 7.8±0.8 vs. Aged: 12.1±1.5 RFU/cell; *P*=0.023). The presence of antimycin A did not alter CellROX oxidation in control or aged skeletal muscle cells (see Figure 1F), suggesting that CellROX oxidation reflects mainly cytosolic, and not mitochondrial-derived ROS. Together, these findings suggest altered metabolic pathways may be required for the provision of energy in aged myotubes because of mitochondrial dysfunction.

### Replicative ageing is associated with altered expression of genes related to energy metabolism

Because replicative ageing was associated with altered mitochondrial function and increased production of mitochondrial superoxide, the mRNA expression of key mitochondrial and antioxidant-related genes was quantified. There was no significant main effect of age or time on drp1 expression in myotubes (Figure 2A). A significant main effect of time (*P*=0.005) was found on mfn2 expression in myotubes. Multiple comparisons revealed mfn2 expression was 2-fold higher in aged myotubes over 48 h versus control (*P*=0.024). Although there was a main effect of age (*P*=0.045) observed on Parkin expression in myotubes, multiple comparisons revealed no significant differences in Parkin expression between control and aged myotubes, regardless of the timepoint. A significant main effect of age (*P*=0.015) was discovered for PGC-1α expression, and PGC-1α expression was 3-fold higher in control myotubes over aged at 24 h (*P*=0.027). There was a significant main effect of age on sirt1 expression in myotubes (*P*=0.006). Over 48 h, sirt1 expression was 2.1-fold higher in aged myotubes versus control (*P*=0.003). A main effect of age was found on Tfam expression in myotubes (*P*=0.012). Tfam expression was 2.1-fold higher in aged myotubes compared with control (*P*=0.013). Overall, transcriptional profiles of aged myotubes suggest reduced mitochondrial biogenesis.

**Figure 2.**
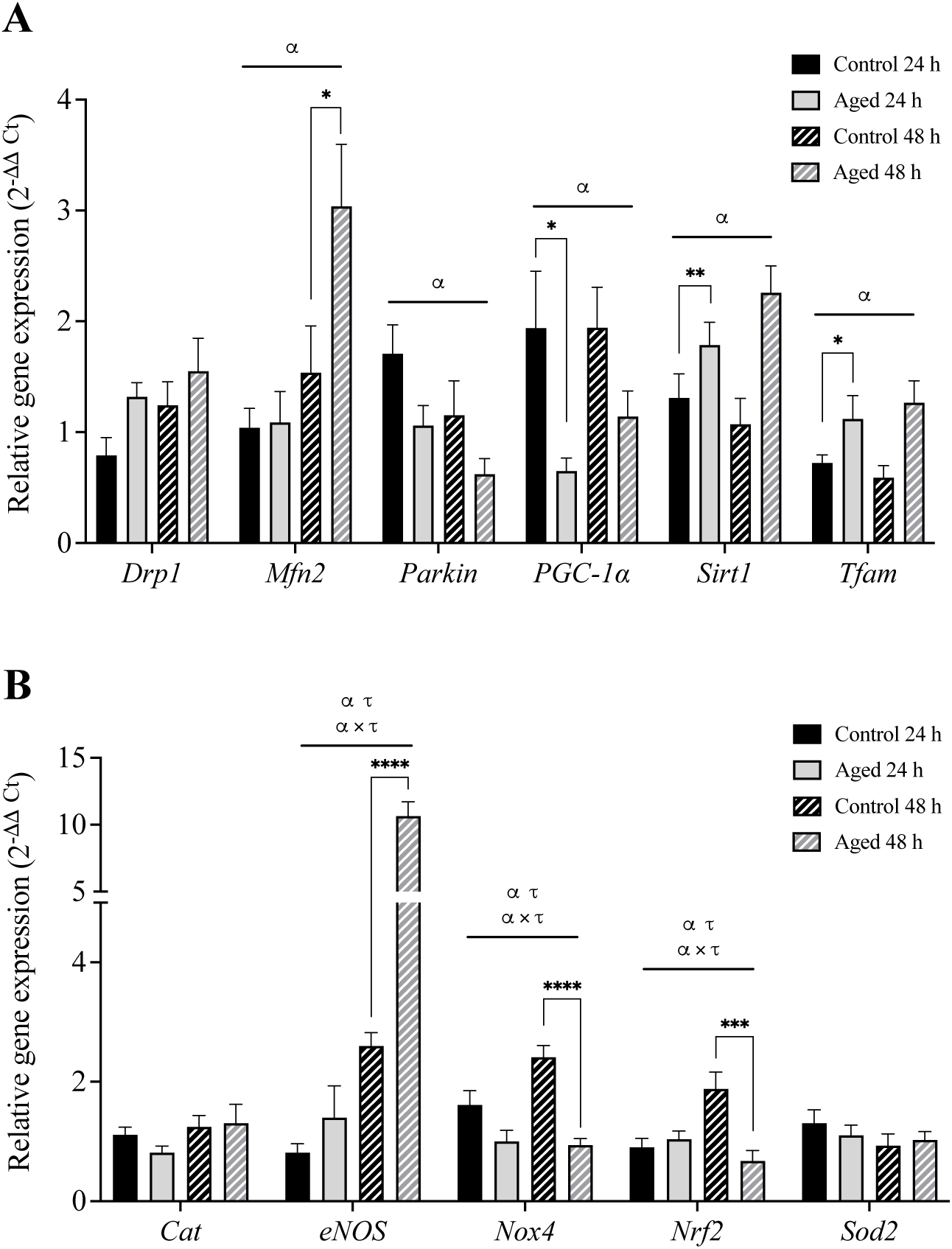
Replicative ageing alters the expression of mitochondrial and antioxidant-related genes. (A)Mitochondrial and (B) antioxidant related genes. C2C12 myotubes were lysed over 72-120 h for analysis of gene expression. Data are means±SEM from 3 independent experiments run in duplicate. Statistical significance was determined by a two-way ANOVA, with age and time as factors. Multiple comparisons performed by Sidak’s test to determine differences in gene expression between ages within each time point. ^α^ significant main effect of age, ^τ^ significant main effect of time (*P*<0.05). \**P*<0.05, \*\**P*<0.01, \*\*\**P*<0.001 and \*\*\*\**P*<0.0001. Control and aged myotubes are denoted by solid black and grey bars, respectively.

There was no main effect of age or time on catalase and sod2 expression in myotubes (see Figure 2B). Significant main effects of age (*P*<0.001) and time (*P*<0.001) were discovered for eNOS expression in myotubes, and an age × time interaction (*P*<0.001). Multiple comparisons revealed eNOS expression was 4.1-fold higher in aged myotubes over 48 h versus control (*P*<0.001). There was a main effect of age (*P*<0.001) and time (*P*=0.047) for nox4 expression in myotubes, and an age × time interaction (*P*=0.025). Nox4 expression increased 2.6-fold over 48 h in control myotubes versus aged (*P*<0.001). A main effect of age (*P*=0.049) and time (*P*=0.048) was found on nrf2 expression in myotubes (Figure 2), as well as an age × time interaction (*P*<0.001). Nrf2 expression was increased 2.8-fold in control versus aged myotubes over 48 h (*P*<0.001). Together, these findings are suggestive of reduced NO bioavailability and lowered induction of ROS sensitive transcripts in aged myotubes.

### Replicative ageing alters the metabolic signatures of C_2_C_12_ myotubes

To explore whether respiratory and transcriptional changes associated with replicative ageing were reflected at the metabolite level, untargeted metabolomics were performed. Principal component analysis (PCA) was performed to observe major variances in the data. PCA scores plot of PC1 (48.27%) against PC2 (28.82%) revealed moderate clustering of age in terms of separation (Figure 3A) and together explained a cumulative variance of 77.09%. Meanwhile a total of 6 components were required to explain 95% of variance in the data. The overall metabolic profile of myotubes suggested: 1) control and aged myotubes are metabolically distinct; 2) aged myotubes display similar variance to control despite one distinct sample in the aged group.

**Figure 3.**
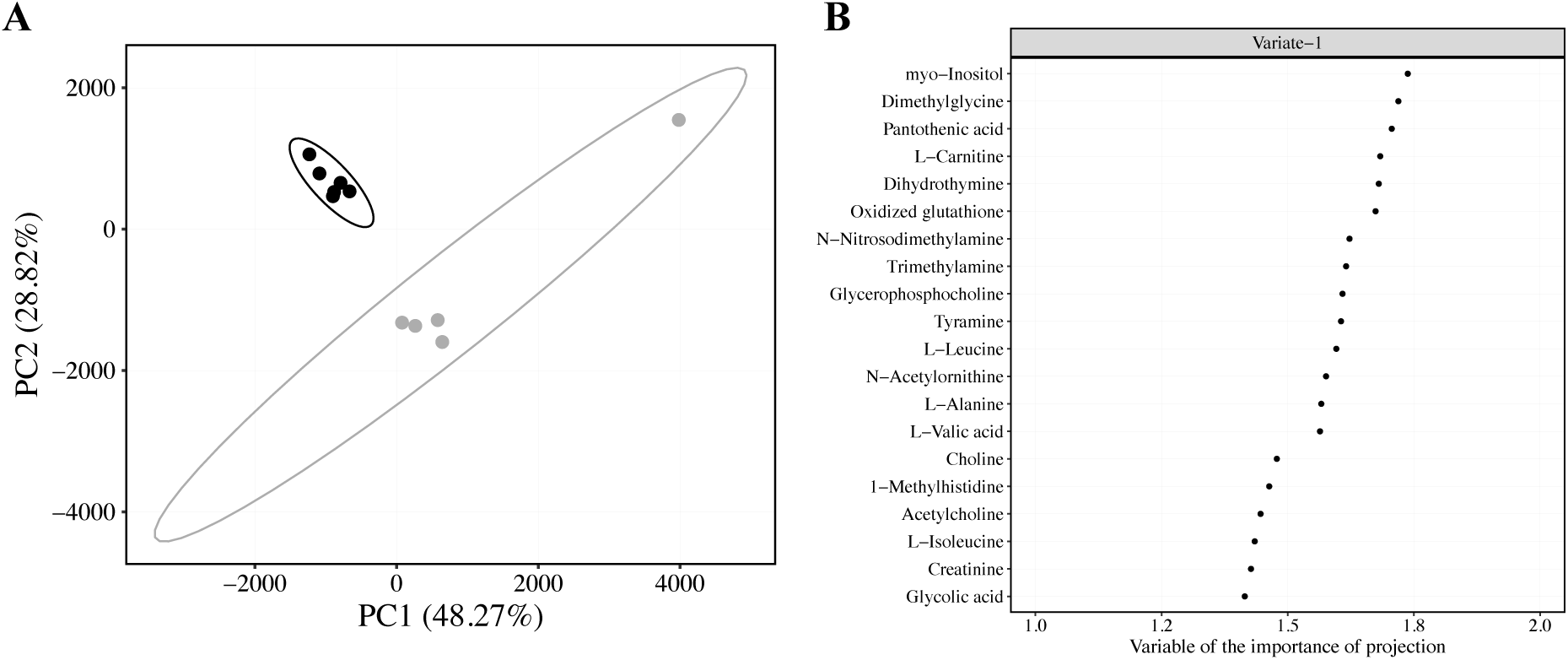
Multivariate analyses of control and aged myotube metabolomes. A) Principal component analysis scores of control and aged myotubes, coloured by age (control = black circles, n=6 and aged = grey circles, n=5). Brackets report the variance explained by the PC. Six PCs were required to achieve 95% explained variance. Only PC1 and PC2 are show in the Figure for clarity. Ellipses represent 95% confidence region. B) VIP scores of PLS-DA model (ROC = 1) built on age-dependent differences in myotubes. A lower threshold of 1 was used on latent variable one to select metabolites from the model. The top 20 representative metabolites/bins are presented for clarity.

To identify distinctions between the metabolic profiles of control and aged skeletal myotubes, age-related differences were enhanced using a cross-validated PLS-DA model. Optimal model complexity was found to be a single-variate model and variable of the importance of projection (VIP) scores were subsequently extracted (Figure 3B). The metabolite level comparison of control and aged myotubes (Figure 4) revealed that, Pantothenic acid, L-Carnitine, Glycerophosphocholine, L-Leucine, N-Acetylornithine, L-Alanine, 1-Methylhistidine, L-Isoleucine, Taurine, L-Tryptophan, cis-Aconitic acid/L-Acetylcarnitine and N,N-Dimethylformamide were significantly lower in aged versus control skeletal myotubes (see Figure 4). On the other hand, myo-Inositol, Dimethylglycine, Oxidized glutathione, Trimethylamine, Tyramine, Choline, Glycolic acid, L-Valine, L-Tyrosine, N-Alpha-acetyllysine and Pyruvic acid were significantly higher in aged versus control skeletal myotubes.

**Figure 4.**
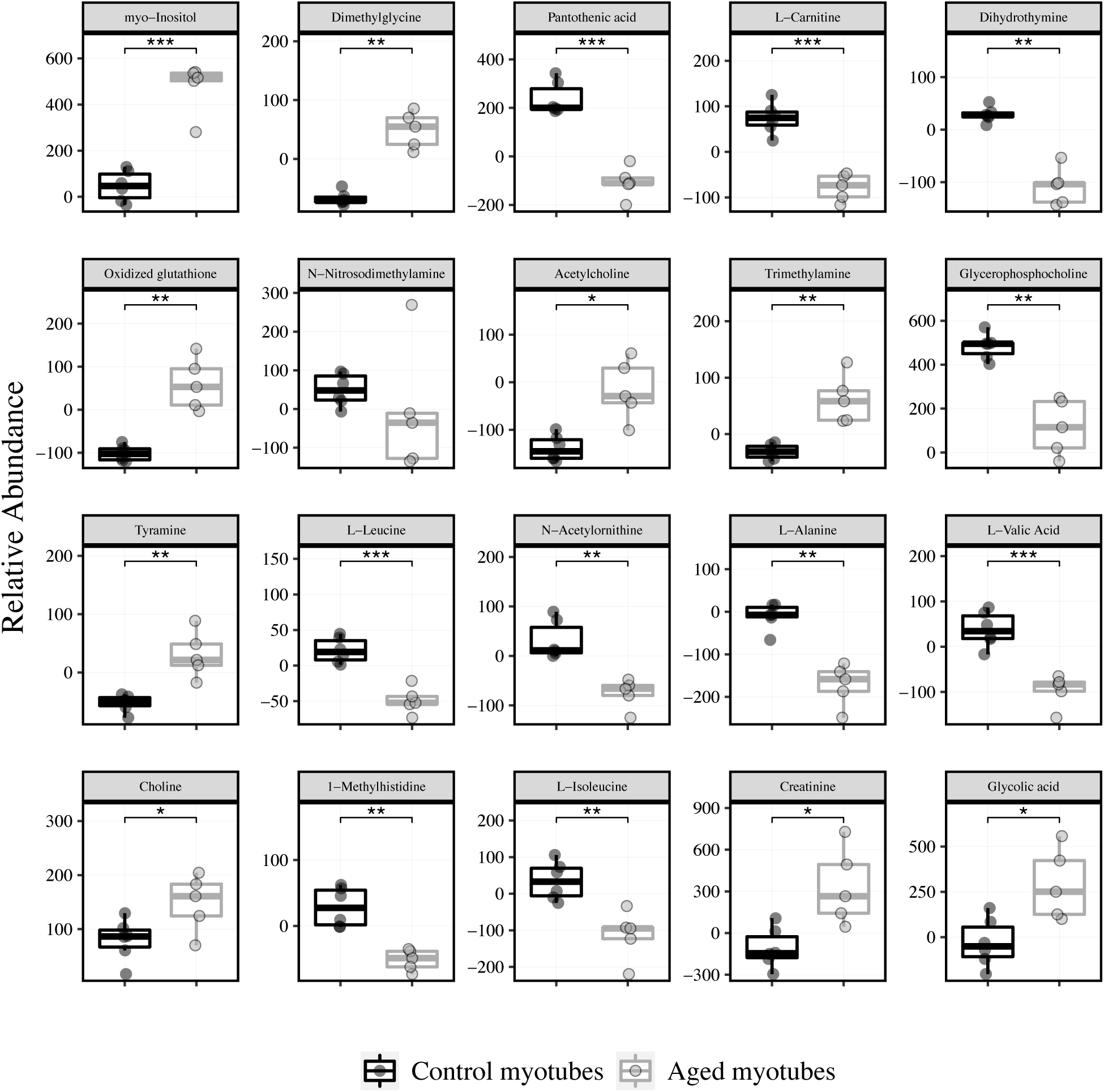
Boxplots of individual metabolites (VIP>1) in control and aged myotubes. Control (black outline, n=6) and aged (grey outline, n=5) myotubes. *, **, *** and **** represent false discovery rate adjusted *P*-value less than 0.05, 0.01, 0.001 and 0.0001, respectively.

Metabolite set enrichment analysis (MSEA) was performed to extract further metabolic pathway level information. Supplementary Table 2 summarises all metabolites selected for MSEA. Three significantly over-represented pathways were identified for control and aged skeletal myotubes (see Supplementary Table 3). Out of the three pathways, metabolites present in two pathways including aminoacyl-tRNA biosynthesis and valine, leucine and isoleucine biosynthesis were predominately lower overall in aged versus control skeletal myotubes.

## Discussion

Replicative ageing perturbs the regenerative potential of human and murine skeletal muscle myoblasts, but it is unclear whether replicative ageing impacts skeletal muscle cellular energy metabolism. Here, we used a combination of plate-based respirometry and untargeted metabolomics to demonstrate that replicative ageing perturbs mitochondrial function and alters energy metabolism of murine skeletal myotubes. Our findings implicate bioenergetic uncoupling, increased superoxide production and down-regulated amino acid metabolism as features of replicative ageing that may exacerbate the age-related impairments to skeletal muscle function.

Mitochondria are essential for skeletal muscle function via energy production and intracellular signalling. Using plate-based respirometry, we observed reduced coupling efficiency, lower maximal respiratory capacity, and lower reliance on oxidative phosphorylation for ATP synthesis in replicatively aged myotubes. Reductions in the proportion of oxygen consumption utilized for ATP synthesis are indicative of increased proton leak. However, one previous study found no evidence for increased (adenine nucleotide translocase-mediated) proton leak with replicative ageing of young and older human-derived skeletal myoblasts (Pääsuke et al., 2016), although uncoupling of skeletal muscle mitochondria has been reported in older adults (Porter et al., 2015). Further studies are necessary to understand the potential source of age-related muscle mitochondrial proton leaks.

The relative contribution of oxidative phosphorylation to total ATP synthesis declined with replicative ageing in this study, which was met by an increase in glycolysis to meet energy demands. This finding supports previous work demonstrating increased reliance on glycolysis in cultured myotubes obtained from elderly adults (Marrone et al., 2018). Together, these findings suggest that replicative ageing results in changes to cellular energy metabolism. Another observation of this study was that replicative ageing was associated with diminished maximal respiratory capacity, which implies impaired maximal electron transfer capacity. However, this finding should be interpreted with caution. We cannot exclude the possibility that the maximal respiratory response obtained was not optimal – owed to an ATP crisis induced by oligomycin – though we employed identical concentrations of oligomycin and FCCP to control and aged myotubes. Overall, replicatively aged skeletal myotubes recapitulate some features of human skeletal muscle associated with mitochondrial dysfunction (Marcinek et al., 2005; Porter et al., 2015; Tonkonogi et al., 2003).

Ageing muscle has repeatedly been linked with increased oxidative stress, where the production of oxidants exceeds the intracellular antioxidant capacity (Chabi et al., 2008; Holloway et al., 2018; Vasilaki et al., 2006). Here, we report that replicative ageing is associated with increased production of mitochondrial superoxide. Similarly, one previous study has documented elevated levels of mitochondrial superoxide in elderly myotubes cultured in the presence of N-acetyl cysteine or high glucose (Marrone et al., 2018), although the increase in superoxide was not observed in the absence of cell stressors. The increase in myotube ROS production with replicative ageing was not limited to the mitochondrial compartment. We also observed that increased ROS production in aged myotubes was not exclusive to mitochondria, suggesting that age-related increases in ROS production are both mitochondrial and cytosolic in origin. To this point, previous studies have demonstrated age-related increases in ROS production from mitochondria (Vasilaki et al., 2006) and extramitochondrial enzymes such as the NADPH oxidases (Pearson et al., 2014; Sullivan-Gunn & Lewandowski, 2013). Interestingly, our gene expression data did not indicate a robust upregulation of antioxidant related transcripts in aged myotubes, despite increased rates of ROS production.

Another key finding of our study was that replicative ageing culminated in changes to the expression of genes associated with energy metabolism. The abundance of mfn2 was acutely upregulated in aged myotubes, which lends support to studies demonstrating an increased ratio of fusion:fission related proteins in ageing skeletal muscle (Joseph et al., 2013; Leduc-Gaudet et al., 2015; Mercken et al., 2017). Similarly, aged myotubes presented increased mRNA expression of Tfam, which serves to increase the synthesis of electron transfer system complex subunits. Some (Lezza et al., 2001), but not all (Welle et al., 2003) studies have also reported increased expression of Tfam in older skeletal muscle tissue. Despite increased Tfam expression, aged myotubes had less abundant PGC-1α and NRF2 mRNA, which implies reduced capacity for mitochondrial biogenesis with replicative ageing. Similar results have been documented at the transcriptional level in ageing skeletal muscle *in vivo* (Ghosh et al., 2011; Shavlakadze et al., 2019; Su et al., 2015), suggesting aged myotubes are less sensitive to biochemical queues that initiate mitochondrial biogenesis (Ljubicic et al., 2009; Ljubicic & Hood, 2009).

To further characterize the metabolic impact of replicative ageing, cellular metabolites were quantified by untargeted metabolomics. Overall, the metabolic signatures of control and aged myotubes were profoundly different, which was largely due to lower levels of aminoacyl-tRNA and BCAA biosynthesis-related metabolites in aged myotubes. Reductions in the abundance of branched chain amino acids in ageing rodent muscle have been reported previously (Zhuang et al., 2021) and suggest a lowered potential for translation and protein synthesis, with potential implications for the fusion capacity of aged muscle cells (Brown et al., 2017). Furthermore, given that branched chain amino acids can be metabolised and can participate in the TCA cycle (Ye et al., 2020), these metabolites may have been sequestered to support metabolic flux in aged myotubes. Aside from amino acids, L-Carnitine levels were lowered by ageing in this study, implying lowered capacity for fatty acid oxidation. This observation aligns with *in vivo* data where older adults display impaired free fatty acid oxidation and mitochondrial dysfunction (Petersen et al., 2003). Moreover, lowered capacity for oxidation of free fatty acids supports our respirometry data, where replicative ageing was associated with reduced reliance on oxidative phosphorylation for ATP synthesis. Glutathione is a potent cellular antioxidant, and a lower ratio of reduced to oxidized glutathione is indicative of increased oxidative stress. The reported increase in abundance of oxidized glutathione in aged myotubes presented supports studies reporting elevated oxidative stress in aged skeletal muscle cells (Beccafico et al., 2007; Drew et al., 2003; Fulle et al., 2005; Marrone et al., 2018; Mecocci et al., 1999; Vasilaki et al., 2006). This increase in glutathione may be due the accumulation of hydrogen peroxide given the age-related increase in myotube ROS production we observed (Fulle et al., 2005).

This study identified two metabolites that have recently been posited to have novel roles in skeletal muscle cellular ageing. One recent study reported that myo-inositol promoted anti-ageing effects via stimulatory effects on phosphatase and tensin homolog (PTEN)-dependent mitophagy in worm and mouse skeletal muscle (Shi et al., 2020). Here, increased myo-inositol levels were identified in aged myotubes, which may reflect a compensatory mechanism to maintain mitochondrial health. A second metabolite, dihydrothymine, thought to reflect DNA damage, was less abundant in aged myotubes in this study. Interestingly, one recent investigation of human skeletal muscle ageing using metabolomics reported an age-related increase in dihydrothymine (Wilkinson et al., 2020). In contrast, we observed replicatively aged myotubes present lower levels of this metabolite. The discrepancies between the findings could well reflect differences in replicative and chronological ageing, or perhaps the model systems used. Indeed, C_2_C_12_ myoblasts fail to fully recapitulate the transcriptional and metabolic features of human skeletal muscle cells (Abdelmoez et al., 2020) and lack the intra- and extra-cellular environment that is present *in vivo* (Aas et al., 2013).. Either way, further research is necessary to elucidate the potential relationship between dihydrothymine and skeletal muscle ageing.

## Conclusion

To conclude, replicative ageing of skeletal myotubes is associated with mitochondrial uncoupling and altered energy metabolism. Our findings implicate bioenergetic uncoupling, increased ROS production and impaired amino acid metabolism in the development of cellular ageing. With the known roles of mitochondrial dysfunction and altered metabolic signatures in chronological skeletal muscle ageing, the reported impact of replicative ageing on skeletal muscle cellular metabolism might contribute to the progression of sarcopenia and exercise intolerance in older humans.

## Availability of data and material

All the data presented in this manuscript are available upon request. Metabolomics data are deposited at EBI.ac.uk/metabolights/MTBLS2024.

## Authors’ contributions

DGS and CES conceived the study and designed experiments. DGS and JB designed the respiration experiments using the Seahorse XFe96 Analyzer. DGS and MMP collected and analysed metabolomics data. DGS collected and analysed the data, DGS, MMP, JB, RD, HJ, DHJT and CES interpreted the data. DGS and CES wrote the manuscript and all authors revised it critically. All authors provided final approval of the version to be published and agree to be accountable for all aspects of the work in ensuring that questions related to the accuracy or integrity of any part of the work are appropriately investigated and resolved. All people designated as authors qualify for authorship, and all those who qualify for authorship are listed. DGS is the guarantor for the work and/or conduct of the study, had full access to all the data in the study and takes responsibility for the integrity of data and the accuracy of the data analysis, and controlled the decision to publish.

## Acknowledgements

The authors acknowledge use of the Mitochondrial Profiling Centre, a core resource supported by the University of Birmingham, and the High-Field NMR Facility at The University of Liverpool. This study was supported by the BBSRC funded Industrial CASE (iCASE) award (BB/P504385/1).

