## Supplementary Files for "Replicative ageing perturbs the metabolic signature of murine C_2_C_12_ skeletal myotubes"

**Supplementary information**

**Table S1.** Primer sequences for Mus musculus with product length. All primers were used under the same cycling conditions.

| Gene | Accession | Sequence  Forward/Reverse or Anchor nucleotide | Product length (bp) |
| --- | --- | --- | --- |
| Polr2b (RP2β) | NM_153798.2 | F: GGTCAGAAGGGAACTTGTGGTAT  R: GCATCATTAAATGGAGTAGCGTC | 197 |
| CAT | NM_009804 | AN: 324 | 96 |
| SOD2 | NM_013671 | AN: 1769 | 103 |
| Dnm1 (DRP1) | NM_152816 | AN: 1337 | 104 |
| MFN2 | NM_113201 | AN: 1709 | 93 |
| Ppargc1a (PGC-1α) | NM_008904 | AN: 4601 | 122 |
| Sirt1 | NM_001159589.2 | F: ACAATTCCTCCACCTGAG  R: GTAACTTCACAGCATCTTCAA | 124 |
| Tfam | NM_009360.4 | F: TCTTGGGAAGAGCAGATGGC  R: GTCTCCGGATCGTTTCACACT | 72 |
| eNOS | NM_008713.4 | F: GGTTGCAAGGCTGCCAATTT  R: TAACTACCACAGCCGGAGGA | 106 |
| Nox2 | NM_007807.5 | F: CAGAACCAACACTTAACCTT  R: CAACCACACCAGAATGAC | 84 |
| Nox4 | NM_015760.5 | F: TCCCTCCTATGGGCAATGTG  R: TGCACATCAAGCCTGGACAA | 177 |
| Prkn (PARKIN) | NM_001317726.1 | AN: 724 | 92 |
| NRF2 | NM_010902.4 | F: GGACATGGAGCAAGTTTGGC  R: CCAGCGAGGAGATCGATGAG | 164 |

**Table S2.** Metabolites with VIP scores >1 and entered for pathway analysis for comparison of control and aged skeletal myotubes under CTRL conditions.

| HMDB | Metabolite | Bin  [*Overlap] | Direction of change vs. control |
| --- | --- | --- | --- |
| HMDB0000001 | 1-Methylhistidine | 29 | ↓ |
| HMDB0000407 | 2-Hydroxy-3-methylbutyric acid | 355/356 | - |
| HMDB0004096 | 5-Methoxyindoleacetate | 40 | - |
| HMDB0000895 | Acetylcholine | 285/286 | - |
| HMDB0001890 | Acetylcysteine | 210 | - |
| HMDB0000034 | Adenine | 18/20 | - |
| HMDB0000097 | Choline | 186 | ↑ |
| HMDB0000072 | cis-Aconitic acid | 182 | ↓ |
| HMDB0000064 | Creatine | 103 | - |
| HMDB0000562 | Creatinine | 198 | - |
| HMDB0000079 | Dihydrothymine | 336/337 | - |
| HMDB0000092 | Dimethylglycine | 212 | ↑ |
| HMDB0000086 | Glycerophosphocholine | 184 | ↓ |
| HMDB0000115 | Glycolic acid | 101 | ↑ |
| HMDB0001273 | Guanosine triphosphate | 62/69 | - |
| HMDB0000201 | L-Acetylcarnitine | 182 | - |
| HMDB0000161 | L-Alanine | 319 | ↓ |
| HMDB0000062 | L-Carnitine | 266 | ↓ |
| HMDB0000641 | L-Glutamine | 258 | - |
| HMDB0000172 | L-Isoleucine | 359 | ↓ |
| HMDB0000687 | L-Leucine | 308 | ↓ |
| HMDB0000159 | L-Phenylalanine | 36 | - |
| HMDB0000929 | L-Tryptophan | 44 | ↓ |
| HMDB0000158 | L-Tyrosine | 45 | ↑ |
| HMDB0000883 | L-Valine | 354 | ↑ |
| HMDB0000211 | myo-Inositol | 141 | ↑ |
| HMDB0003357 | N-Acetylornithine | 312 | ↓ |
| HMDB0000446 | N-Alpha-acetyllysine | 313/314 | ↑ |
| HMDB0031419 | N-Nitrosodimethylamine | 120/190 | - |
| HMDB0001888 | N,N-Dimethylformamide | 200 | ↓ |
| HMDB0003337 | Oxidized glutathione | 282 | ↑ |
| HMDB0000210 | Pantothenic acid | 360 | ↓ |
| HMDB0000243 | Pyruvic acid | 273 | ↑ |
| HMDB0000251 | Taurine | 159 | ↓ |
| HMDB0000906 | Trimethylamine | 213 | ↑ |
| HMDB0000306 | Tyramine | 209 | ↑ |

↑ significantly higher and ↓ significantly lower in aged vs. control cells (*P*<0.05). – similar between control and aged.

**Table S3.** Pathway analysis results for control and aged skeletal myotubes. Reporting raw & BH adjusted *P* values, number of hits, pathway impact and matches.

| **Pathway** | **Raw *P-*value** | **BH *P-*value** | **Hits** | **Impact** | **Matches** |
| --- | --- | --- | --- | --- | --- |
| Aminoacyl-tRNA biosynthesis | <0.0001 | 0.0006 | 8 | 0 | L-Phenylalanine; L-Glutamine; L-Valine; L-Alanine; L-Leucine; L-Isoleucine; L-Tryptophan; L-Tyrosine; |
| Valine, leucine and isoleucine biosynthesis | 0.0006 | 0.0250 | 3 | 0 | L-Valine; L-Leucine; L-Isoleucine |
| Phenylalanine, tyrosine and tryptophan biosynthesis | 0.0031 | 0.0856 | 2 | 1 | L-Phenylalanine; L-Tyrosine |

Metabolites in red are lower in aged versus control, whereas those in black are similar between control and aged myotubes.
